# Heterogeneous distribution of sex ratio distorters in natural populations of the isopod *Armadillidium vulgare*

**DOI:** 10.1101/2022.09.29.510044

**Authors:** Sylvine Durand, Baptiste Lheraud, Isabelle Giraud, Nicolas Bech, Frédéric Grandjean, Thierry Rigaud, Jean Peccoud, Richard Cordaux

## Abstract

In the isopod *Armadillidium vulgare*, many females produce progenies with female-biased sex ratios, due to two feminizing sex ratio distorters (SRD): *Wolbachia* endosymbionts and the *f* element. We investigated the distribution and population dynamics of these SRD and mitochondrial DNA variation in 16 populations from Europe and Japan. Confirming and extending results from the 1990’s, we found that the SRD are present at variable frequencies in populations, and that the *f* element is overall more frequent than *Wolbachia*. The two SRD never co-occur at high frequency in any population, suggesting an apparent mutual exclusion. We also detected *Wolbachia* or the *f* element in some males, which likely reflects insufficient titer to induce feminization or presence of masculinizing alleles. Our results are consistent with a single integration event of a *Wolbachia* genome in the *A. vulgare* genome at the origin of the *f* element, which contradicts an earlier hypothesis of frequent losses and gains. We identified strong linkage between *Wolbachia* strains and mitochondrial haplotypes, but no association between the *f* element and mitochondrial background. Our results open new perspectives on SRD evolutionary dynamics in *A. vulgare*, the evolution of genetic conflicts and their impact on the variability of sex determination systems.

## 1. Introduction

Sex ratio distorters (SRD) are selfish genetic elements located on sex chromosomes or transmitted by a single sex, which skew the proportion of males and females in progenies towards the sex that enhances their own vertical transmission [1]. Major SRD types include sex chromosome meiotic drivers [2,3], some B chromosomes [4] and selfish cytoplasmic genetic entities [5,6,7]. Collectively, they are found in a wide range of animal and plant species and they have had a tremendous impact on the ecology and evolution of their host species [8,9]. One of the most emblematic SRD is the bacterial endosymbiont *Wolbachia* [10,11]. *Wolbachia* is a cytoplasmic, maternally inherited alpha-proteobacterium found in a wide range of arthropods and nematodes. In arthropods, *Wolbachia* often manipulates host reproduction in favor of infected females, thereby conferring itself a transmission advantage. This is achieved through various strategies, three of which causing sex ratio distortions towards females: male killing, thelytokous parthenogenesis and feminization of genetic males [6,7,10,11].

In the terrestrial isopod *Armadillidium vulgare*, chromosomal sex determination follows female heterogamety (ZZ males and ZW females) [12–14]. However, many females produce progenies with female-biased sex ratios, due to the presence of two feminizing SRD: *Wolbachia* endosymbionts and a locus called the *f* element [6,15,16]. *Wolbachia* symbionts cause ZZ genetic males to develop as phenotypic females [17]. Three *Wolbachia* strains have been described in *A. vulgare*, for which feminization induction has been demonstrated (*w*VulC and *w*VulM strains [18,19]) or is strongly suspected (*w*VulP strain [20]). The *f* element is a nuclear insert of a large portion of a feminizing *Wolbachia* genome in the *A. vulgare* genome [21]. The *f* element induces female development, as a W chromosome does, and it shows non-Mendelian inheritance, making it an SRD [21,22]. These SRD may cause turnovers in sex determination mechanisms [6,15,23] and they could explain why sex chromosome systems are so variable in terrestrial isopods [24–27].

Testing this hypothesis requires characterizing the evolutionary dynamics of SRD such as *Wolbachia* and the *f* element in natural populations. In *A. vulgare*, this characterization is quite limited because prior studies were mostly restricted to a narrow geographic area (western France), sometimes focusing solely on *Wolbachia* [20,28–31]. The only exception is a 1993 study [32], which collated and extended results from the early 1980’s [33,34]. The main observations were that *Wolbachia* and the *f* element are present at variable frequencies in field populations, and the *f* element is more frequent than *Wolbachia*. However, earlier studies were limited by the lack of molecular tests for *Wolbachia* and/or the *f* element, preventing any direct assessment of SRD presence. Instead, the authors used a complex, indirect procedure combining a physiological test and crossings [32]. In addition to being tedious and time-consuming (generation time is one year in this species), this procedure did not allow direct and undisputable assessment of SRD presence. Moreover, it could only be run on females and therefore provided no information on SRD presence in males. Finally, it could not reveal individuals potentially carrying both SRD.

Here, we took advantage of the availability of molecular markers to directly assess SRD presence in males and females from *A. vulgare* field populations from Europe and Japan. This approach allowed us to circumvent the limitations of previous studies, and to revisit the population dynamics of *Wolbachia* and the *f* element in this species and their association mitochondrial lineages.

## 2. Materials and Methods

647 *A. vulgare* individuals from 16 natural populations across Europe and Japan were collected by hand. Individuals were sexed and stored in alcohol or at −20°C prior to DNA extraction. Total genomic DNA was extracted from the head and legs of each individual, as described previously [21]. We used four molecular markers to assess the presence of *Wolbachia* and the *f* element in DNA extracts: *Jtel* [21], *wsp* [35], *recR* [36] and *ftsZ* [37] (Table S1). While *Jtel* is specific to the *f* element, *wsp* and *recR* are specific to *Wolbachia*, and *ftsZ* is present in both the *f* element and *Wolbachia* [21]. We assessed the presence or absence of these markers by PCR, as described previously [21]. Different amplification patterns were expected for individuals with *Wolbachia* only (*Jtel-, wsp+*, *recR+*, *ftsZ+*), the *f* element only (*Jtel+, wsp-, recR-*, *ftsZ+*), both *Wolbachia* and the *f* element present (*Jtel+, wsp+, recR+*, *ftsZ+*) or both *Wolbachia* and the *f* element lacking (*Jtel-, wsp-, recR-*, *ftsZ-*). The few individuals exhibiting other amplification patterns were classified as “undetermined status”. A quantitative-PCR assay was used to measure *Wolbachia* titer in some individuals (see supplementary Methods). To characterize *Wolbachia* strain diversity, *wsp* PCR products were purified and Sanger sequenced using both forward and reverse primers by GenoScreen (Lille, France). Forward and reverse reads were assembled using Geneious^®^ v.7.1.9 to obtain one consensus sequence per individual. To evaluate mitochondrial diversity, we amplified by PCR a ~700 bp-long portion of the Cytochrome Oxidase I (*COI*) gene in all individuals [38]. PCR products were purified and Sanger sequenced as described above. Haplotype network analysis was performed using the *pegas* package [39]. All statistical analyses were performed with R v.3.6.0 [40]. Figures were realized with *ggplot2* [41].

## 3. Results

We tested the presence of *Wolbachia* and the *f* element in 423 females and 224 males from 16 populations across Europe and Japan (Tables 1, S2). While most males lacked both SRD, 48% of females carried at least one of them. The remaining females presumably carry W chromosomes, although the existence of other feminizing elements cannot be formally excluded. As expected for feminizing elements, the SRD were mostly found in females, the *f* element being more frequent than *Wolbachia* overall. Both SRD were found in the same individuals in only 3 females from a single population (Chizé). *Wolbachia*-infected individuals carried one of the three previously known *Wolbachia* strains of *A. vulgare*: *w*VulC (n=62), *w*VulM (n=23) or *w*VulP (n=4).

**Table 1.**
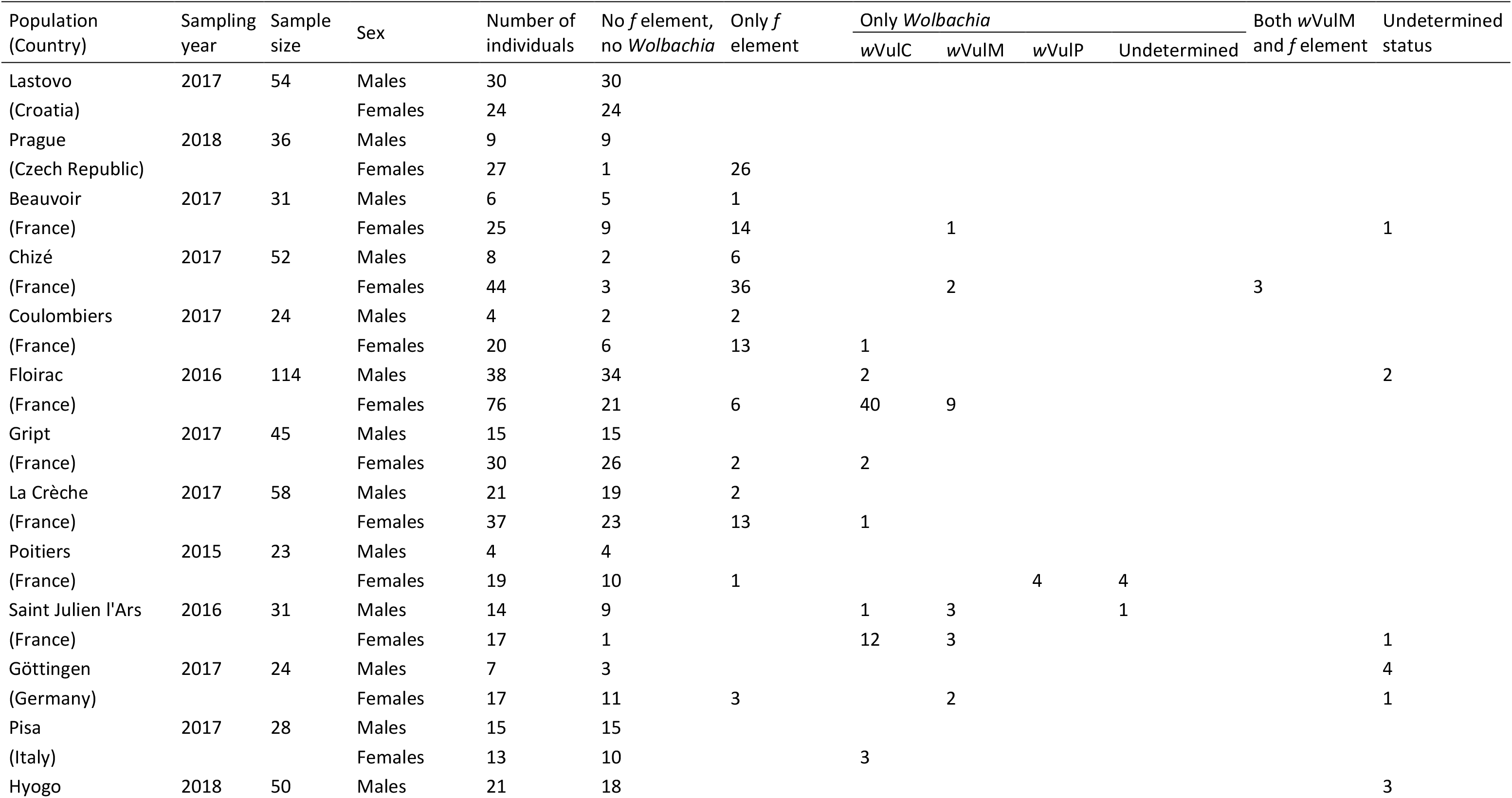
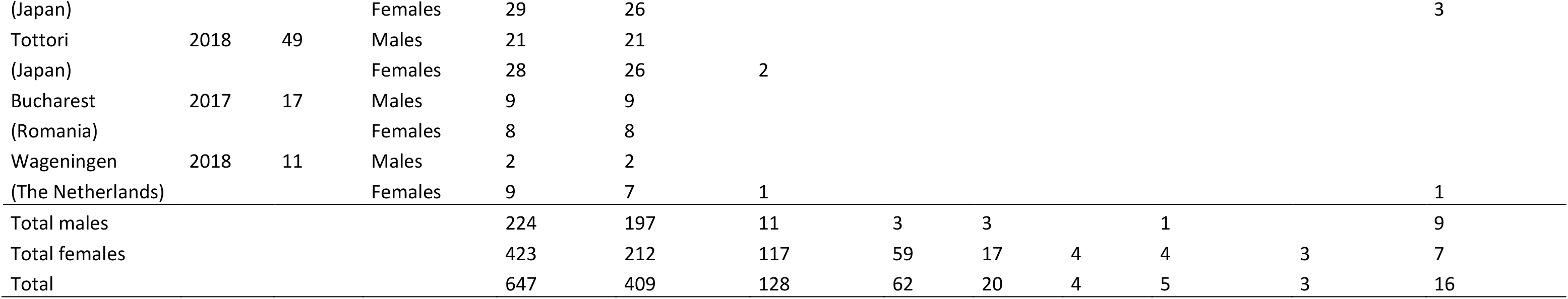
Prevalence of *Wolbachia* and *f* element sex ratio distorters in 16 populations of *Armadillidium vulgare*.

*Wolbachia* and *f* element distribution in females was highly heterogenous among populations (Figure 1a). These SRD were found in 10 and 11 out of 16 populations, but they reached frequencies >10% in only 6 and 7 populations, respectively. The two SRD coexisted in 8 populations. A generalized linear model predicting the frequency of the *f* element as a binomial response by the proportion of individuals carrying *Wolbachia* (each statistical unit being a population) showed that the prevalence of the two SRD was significantly negatively correlated (9.8% of deviance explained, Chi-squared test, *p* < 7.9 × 10^−8^, 14 df) (Figure 1b). Hence, in Floirac, Poitiers, Saint Julien l’Ars and Pisa populations, *Wolbachia* was frequent (23-94% frequency in females) and the *f* element was rare (0-8%). By contrast, the *f* element was frequent (35-96%) and *Wolbachia* was rare (0-11%) in Prague, Beauvoir, Chizé, Coulombiers and La Crèche populations. In the other populations, both SRD were found at low to moderate frequency (0-19%), including 3 populations devoid of both SRD (Lastovo, Hyogo and Bucharest).

**Figure 1.**
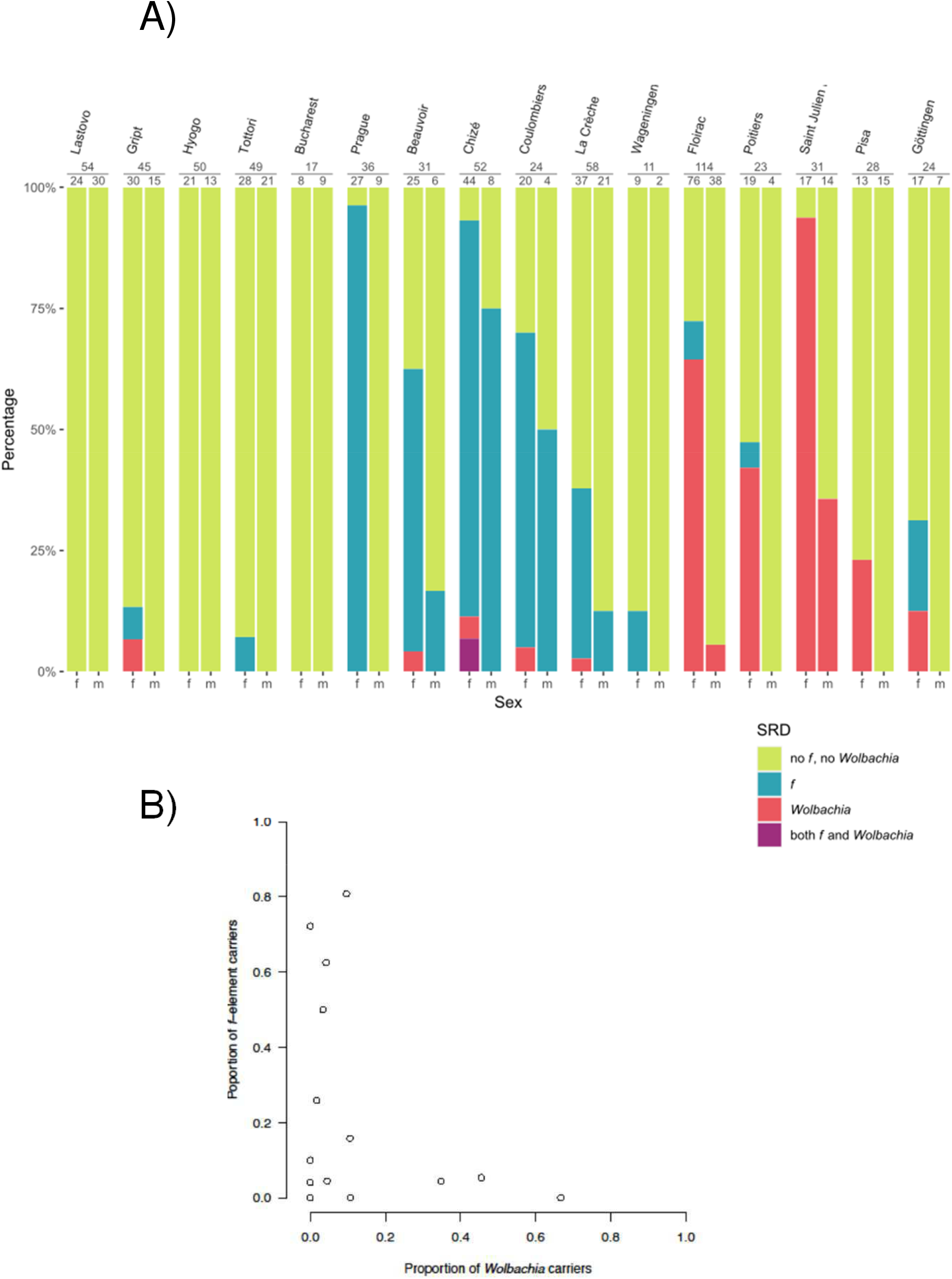
(**A**) Prevalence of *Wolbachia* and the *f* element in males (m) and females (f) from 16 *Armadillidium vulgare* populations. (**B**) Relative proportions of *Wolbachia* and the *f* element in 16 *A. vulgare* populations (represented by open circles).

Males carrying *Wolbachia* or the *f* element were found in 2 and 4 out of 16 populations, respectively. In all cases, these males occurred in populations in which the corresponding SRD were the most prevalent ones in females: Beauvoir, Chizé, Coulombiers and La Crèche for the *f* element, and Floirac and Saint Julien l’Ars for *Wolbachia*. Overall, these males had much lower *Wolbachia* titer than females from their respective populations (Figure S1, Table S3).

The 642 individuals sequenced at the *COI* gene presented a total of 92 segregating sites defining 23 haplotypes (named I to XXIII; GenBank accession numbers in Table S4), with 1 to 7 haplotypes per population (Table S2, Figure 2). The most frequent and widespread haplotype (I) was found in 188 individuals from 10 populations. The second most frequent and widespread haplotype (V) was found in 106 individuals from 7 populations. We found 21 out of the 23 haplotypes among individuals lacking both *Wolbachia* and the *f* element (Table 2, Figure 2). Among individuals carrying the *f* element, 6 haplotypes were found, all but one (I, II, III, V and VI) being shared with individuals lacking both *Wolbachia* and the *f* element, and one (IV) being carried by a single individual in the entire dataset. Among *Wolbachia*-infected individuals, all those carrying *w*VulC were associated with either haplotype V or its close relatives (XI and XII). All individuals carrying *w*VulM were associated with haplotype II and those carrying *w*VulP with haplotype VII. Of the 5 haplotypes found in *Wolbachia*-infected individuals, 4 were shared with individuals lacking both *Wolbachia* and the *f* element (II, V, VII and XII), 2 of which were also shared with individuals carrying the *f* element (II and V), and one (XI) was present in a single individual in the entire dataset.

**Figure 2.**
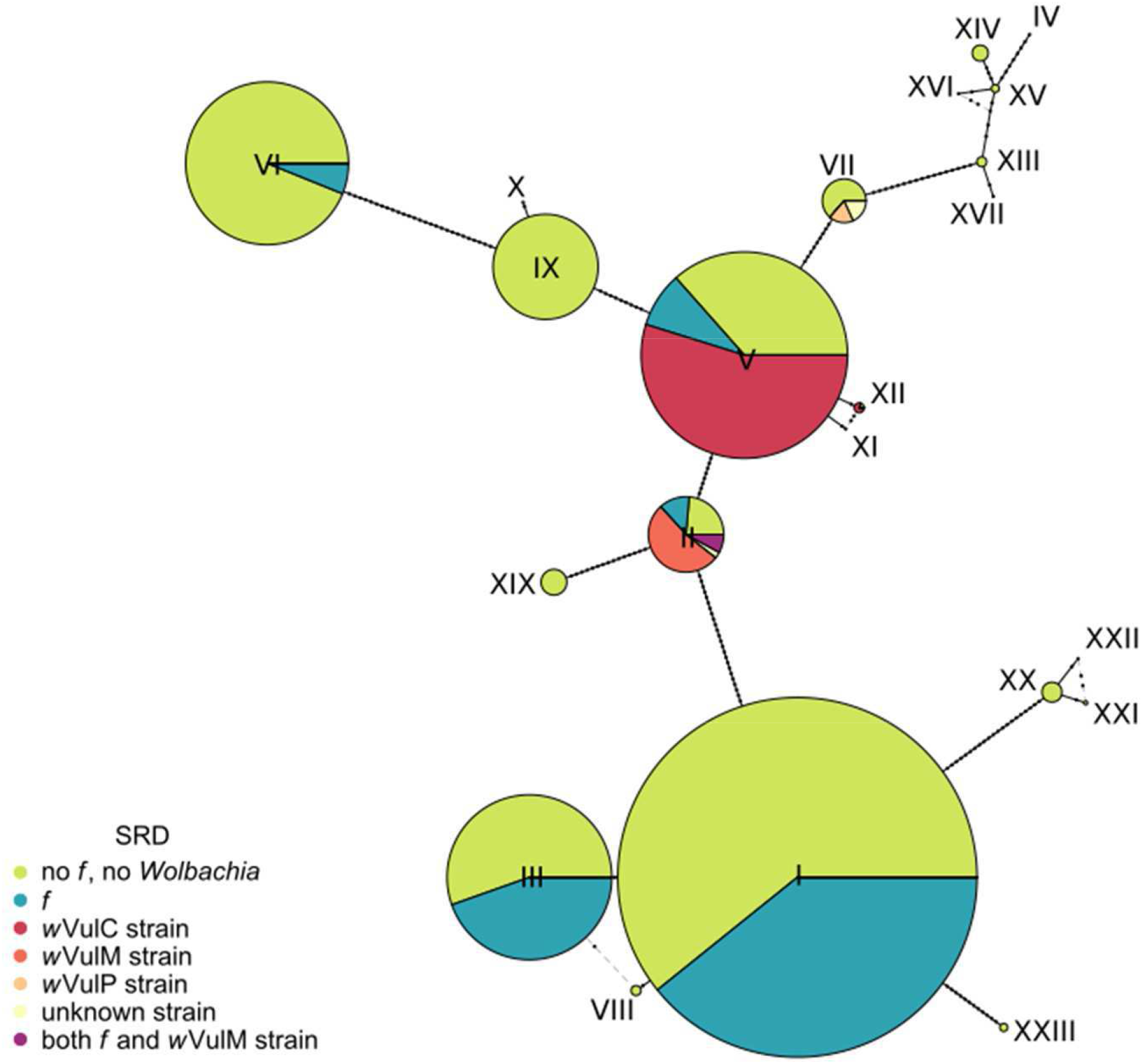
Haplotype network of 23 mitochondrial variants (I-XXIII) from 16 *Armadillidium vulgare* populations. Each circle represents one haplotype and circle diameter is proportional to the number of individuals carrying the haplotype. Branch lengths connecting circles are proportional to divergence between haplotypes. Sex ratio distorter frequencies are color-coded for each haplotype.

**Table 2.** Distribution of mitochondrial haplotypes in 642 *Armadillidium vulgare* individuals from 16 populations.

| Sex ratio distorter status | Number of individuals | Haplotype number | Haplotype list |
| --- | --- | --- | --- |
| No *f* element, no *Wolbachia* | 404 | 21 | I, II, III, V, VI, VII, VIII, IX, X, XII, XIII, XIV, XV, XVI, XVII, XVIII, XIX, XX, XXI, XXII, XXIII |
| *f* element only | 128 | 6 | I, II, III, IV, V, VI |
| *Wolbachia* (wVulC strain) only | 62 | 3 | V, XI, XII |
| *Wolbachia* (wVulM strain) only | 20 | 1 | II |
| *Wolbachia* (wVulP strain) only | 4 | 1 | VII |
| *Wolbachia* (undetermined strain) only | 5 | 2 | II, VII |
| Both wVulM and *f* element | 3 | 1 | II |
| Undetermined status | 16 | 4 | I, V, VI, XIX |

## 4. Discussion

Our results provide direct evidence that the *f* element is overall more frequent than *Wolbachia* in the sampled *A. vulgare* populations. We detected the *f* element in 11 *A. vulgare* natural populations from 4 European countries (Czech Republic, France, Germany and The Netherlands) and Japan. Together with its previous detection in Denmark [21], our results indicate that the *f* element has spread to a wide geographical range. The relative frequencies of the *f* element and *Wolbachia* are highly variable among populations and, in general, when one SRD is frequent, the other SRD is rare. Overall, these results are consistent with earlier results from the 1990’s [32], although no molecular assay allowing direct testing was available at that time and SRD presence or absence was inferred indirectly.

As the *Jtel* marker is located across the site of integration of the *f* element in the *A. vulgare* chromosome [21], *f* element presence in various populations can be explained by a single event of integration of a *Wolbachia* genome in the *A. vulgare* genome. A less parsimonious scenario would require independent insertions at the same chromosomal site, which is highly unlikely. The former scenario contradicts an earlier hypothesis on the evolutionary dynamics of the *f* element, which suggested that the *f* element was unstably integrated in the *A. vulgare* genome, experiencing frequent loss from oocytes and recurrent gain from *Wolbachia* endosymbionts [22,23,42–44]. Under this scenario, multiple independent *f*-like elements would be expected to segregate at low frequencies in populations, they should be integrated in different genomic locations and they should all be able to induce feminization [16]. While our results do not formally invalidate the possibility of additional *f*-like integrations in *A. vulgare* populations, which the *Jtel* marker would not detect, it does not appear to be the most parsimonious hypothesis. Examination of sex ratios from progenies of wild-caught females lacking both SRD may offer further insight into this issue.

Using molecular assays allowed us to circumvent two limitations of the previously used physiological test: the impossibility to detect *Wolbachia* and the *f* element in males, and the impossibility to detect individuals carrying both SRD. Regarding *Wolbachia* presence in males, the historic protocol was only applicable to females per design [29,30,32] and subsequent PCR screens for *Wolbachia* infection have mostly focused on testing females [20,30,31]. In fact, males have seldom been tested and found to carry *Wolbachia* [45]. Here, we detected *Wolbachia* in 7 males from 2 populations (Floirac and Saint Julien l’Ars), carrying either *w*VulC or *w*VulM strains. The failure of feminization by *Wolbachia* most certainly reflects insufficient bacterial titer to induce feminization (Figure S1). These field observations hence support the view that titer is an important factor for successful feminization, as low titer is linked to incomplete feminization and intersexual phenotypes [42,46].

We also detected the presence of the *f* element in 11 males from 4 populations. Historically, the presence of the *f* element in males has been indirectly inferred from crossings and the resulting sex-ratios biases of progenies [22,43,47]. Our results constitute the first direct evidence for the presence of the *f* element in *A. vulgare* males. In all 4 populations in which *f*-carrying males were found, the *f* element was also frequent in females. Altogether, these observations suggest that the 11 males carrying the *f* element also carry the masculinizing dominant *M* allele [16,43,47]. Indeed, the *M* allele is able to restore a male phenotype in individuals carrying the *f* element [16,43,47]. Because of sex ratio selection, the *M* allele is thought to have been selected to restore males in response to female-biased sex ratios caused by the *f* element [47]. Thus, the *M* allele is expected to rise in frequency when the *f* element is frequent in a population [47], which is consistent with our observations. Unfortunately, no molecular marker of the *M* allele is currently available, which prevents any direct assessment of its actual presence in these populations. Thus, we cannot exclude that males carrying the *f* element simply carry non-feminizing variants of this SRD.

Our results show that *Wolbachia* and the *f* element never co-occur at high frequency in any population. This apparent mutual exclusion can be explained considering that co-occurrence of multiple feminizing factors in a population should favor the most transmitted one [16,48]. Hence, *Wolbachia* is expected to lead to the loss of nuclear feminizing elements in *A. vulgare* populations. This situation does not result from an interference between chromosomes and *Wolbachia* within individuals, but from counter selection of nuclear feminizing alleles in a population that becomes increasingly biased towards females. Hence, the rise of *Wolbachia* would associate with the decline of the *f* element in a population. Why, under these circumstances, *Wolbachia* has not invaded all *A. vulgare* populations is still unclear and may reflect fitness effects or possible resistance genes.

As a result, only very few individuals were found to carry both *Wolbachia* and the *f* element. They represent only 3 females, all from the Chizé population (Figure 1a). These were likely born from mothers carrying *Wolbachia* and fathers carrying the *f* element, which are frequent at Chizé. The apparent absence of carriers of both SRD in other populations where these SRD are present could simply be explained by the paucity of males carrying the *f* element.

Mapping SRD distribution onto mitochondrial genealogy showed excellent congruence between *Wolbachia* strains and mitochondrial haplotypes (*w*VulC-V, *w*VulM-II and *w*VulP-VII). Such strong association has previously been noted in *A. vulgare*-*Wolbachia* interactions at a smaller geographic scale [30,31] and, more generally, in many arthropod-*Wolbachia* interactions [49]. This result corroborates the rarity of non-maternal transmission of *Wolbachia* in *A. vulgare*. By contrast, the *f* element was found in 6 different mitochondrial backgrounds (I-VI) scattered across the mitochondrial phylogeny, indicating no particular association between the *f* element and mitochondria. This result confirms and extends earlier data focused on western France and in which *f* element presence in females was indirectly inferred based on sex ratios of their progenies [30]. This observation can be explained by the occasional paternal transmission of the *f* element, which breaks its association with mitochondrial background [16,22,30].

## Data accessibility

All data are provided in the electronic supplementary material.

## Acknowledgments

We thank Drs. Nicolas Cerveau, Rémi Elliautout, Misel Jelic, Shigenori Karazawa, Giuseppe Montesanto and Eveline Verhulst for providing samples.

