## Supplementary material for "Heterogeneous distribution of sex ratio distorters in natural populations of the isopod *Armadillidium vulgare*": Figure S1, Tables S1 to S4

Content of the file: Supplementary Methods, Figure S1 and Tables S1-S4.

**Supplementary Methods**

As several males were found to carry *Wolbachia*, we compared *Wolbachia* titer in the DNA extracts from these males to that measured in females from the same populations. We used a quantitative-PCR assay with dual-labelled probes to estimate the concentration of a *Wolbachia* locus relative to the concentration of a nuclear reference gene that has a single copy per haploid *A. vulgare* genome. The *Wolbachia* locus was amplified by primers R_wol_f (5'-GTTGGGTTCTTGTGTGAAAGC-3'), F_wol_f (5'-TGCTCCACCACTCGTATCAATC-3') and revealed with HEX-labelled fluorescent probe_wol_f (5'-AGCTCAGCCATATCCATCAGTGCA-3'). The nuclear gene (encoding mitochondrial tRNA-leucine ligase) was amplified with primers TleuF (5'-TGT-ACA-CAT-CGA-GCA-GCA-AG3-3'), TleuR (5'-GTG-GCA-CCA-TAA-GAC-TTT-GAG-C-3') and revealed with FAM-labelled probe TleuP (5'-ACG-AAG-TTC-GCC-CTG-TTC-TGC-A-3'). The probes allowed the amplification of both loci in the same reaction using the LightCycler-480 instrument.

**Figure S1. Dose of *Wolbachia* DNA relative to nuclear DNA from males and females from two populations for 20 females and 7 males.** The difference in dose is supported by a Mann–Whitney U test (W = 128, p = 0.000554). *Wolbachia* DNA was detected before the 30th cycle of each quantitative PCR assay. The two males showing the highest relative dose were analysed through DNA extracted from a leg (no other DNA sample was available from these males). For other males, whole-body DNA was used. The median relative dose of males is 0.05920 and that of females is 2.297. The underlying data for this figure can be found in Table S3.

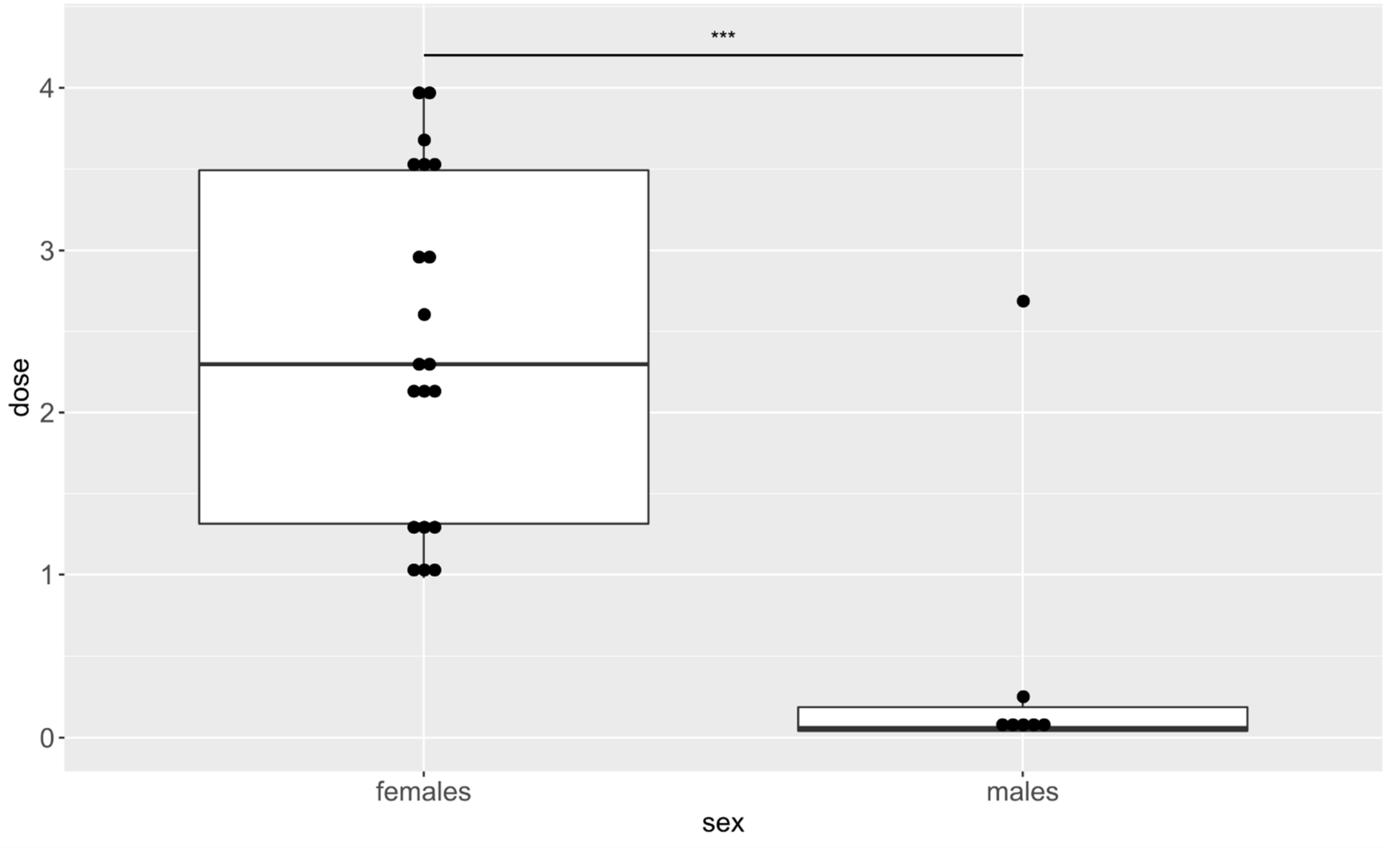

**Table S1. Molecular markers used in this study.**

| Marker name | Target | Primer name | Primer sequence 5'-3' | Tm (°C) | PCR product size (bp) | Reference |
|---|---|---|---|---|---|---|
| *COI* | mitochondria | LCO | GGTCAACAAATCATTAAAGATATTGG | 52 | 700 | [38] |
| | | HCO | TAAACTTCAGGGTGACCAAAAAATCA | | | |
| *Jtel* | *f* element | SubF1 | ACGAAAAGCCGACGTAAAATATTT | 55 | 700 | [21] |
| | | JTelR2 | GAAATAAAAGAGCCTGACT | | | |
| *wsp* | *Wolbachia* | 81F | TGGTCCAATAAGTGATGAAGAAAC | 55 | 650 | [35] |
| | | 691R | AAAAATTAAACGCTACTCCA | | | |
| *recR* | *Wolbachia* | RecR_F_B2 | TGCTTTTCTTCATTTGTTTCCA | 60 | 876 | [36] |
| | | RecR_R_B2 | TTTCTAGGCTGAAGTATGCCACT | | | |
| *ftsZ* | *Wolbachia* and *f* element | ftsZf1 | GTTGTCGCAAATACCGATGC | 47 | 1000 | [37] |
| | | ftsZr1 | CTTAAGTAAGCTGGTATATC | | | |

**Table S2. Information on the 647 *Armadillidium vulgare* individuals from 16 populations included in this study.**

| Country | Population | Individual ID | Phenotypic sex | Presence of *f* element and/or *Wolbachia* | *Wolbachia* strain | CO1 haplotype |
|---|---|---|---|---|---|---|
| Japan | Hyogo | JAP-HYf01 | female | none | n/a | I |
| Japan | Hyogo | JAP-HYf02 | female | none | n/a | VI |
| Japan | Hyogo | JAP-HYf03 | female | none | n/a | I |
| Japan | Hyogo | JAP-HYf04 | female | none | n/a | VI |
| Japan | Hyogo | JAP-HYf05 | female | none | n/a | VI |
| Japan | Hyogo | JAP-HYf06 | female | none | n/a | VI |
| Japan | Hyogo | JAP-HYf07 | female | none | n/a | XIX |
| Japan | Hyogo | JAP-HYf08 | female | none | n/a | VI |
| Japan | Hyogo | JAP-HYf09 | female | none | n/a | XIV |
| Japan | Hyogo | JAP-HYf10 | female | none | n/a | VI |
| Japan | Hyogo | JAP-HYf11 | female | none | n/a | I |
| Japan | Hyogo | JAP-HYf12 | female | none | n/a | VI |
| Japan | Hyogo | JAP-HYf13 | female | none | n/a | XIX |
| Japan | Hyogo | JAP-HYf15 | female | none | n/a | VI |
| Japan | Hyogo | JAP-HYf16 | female | none | n/a | VI |
| Japan | Hyogo | JAP-HYf17 | female | none | n/a | VI |
| Japan | Hyogo | JAP-HYf18 | female | undetermined | n/a | VI |
| Japan | Hyogo | JAP-HYf19 | female | none | n/a | VI |
| Japan | Hyogo | JAP-HYf20 | female | none | n/a | VI |
| Japan | Hyogo | JAP-HYf21 | female | none | n/a | XIX |
| Japan | Hyogo | JAP-HYf22 | female | none | n/a | VI |
| Japan | Hyogo | JAP-HYf23 | female | none | n/a | VI |
| Japan | Hyogo | JAP-HYf24 | female | none | n/a | XIX |
| Japan | Hyogo | JAP-HYf25 | female | undetermined | n/a | VI |
| Japan | Hyogo | JAP-HYf26 | female | undetermined | n/a | XIX |
| Japan | Hyogo | JAP-HYf27 | female | none | n/a | XIX |
| Japan | Hyogo | JAP-HYf28 | female | none | n/a | VI |
| Japan | Hyogo | JAP-HYf29 | female | none | n/a | XIX |
| Japan | Hyogo | JAP-HYf30 | female | none | n/a | VI |
| Japan | Hyogo | JAP-HYm01 | male | none | n/a | XIX |
| Japan | Hyogo | JAP-HYm02 | male | none | n/a | VI |
| Japan | Hyogo | JAP-HYm03 | male | none | n/a | VI |
| Japan | Hyogo | JAP-HYm04 | male | undetermined | n/a | VI |
| Japan | Hyogo | JAP-HYm05 | male | none | n/a | VI |
| Japan | Hyogo | JAP-HYm06 | male | undetermined | n/a | VI |
| Japan | Hyogo | JAP-HYm11 | male | none | n/a | VI |
| Japan | Hyogo | JAP-HYm12 | male | none | n/a | VI |
| Japan | Hyogo | JAP-HYm14 | male | none | n/a | XIX |
| Japan | Hyogo | JAP-HYm16 | male | none | n/a | XIX |
| Japan | Hyogo | JAP-HYm18 | male | none | n/a | XIX |
| Japan | Hyogo | JAP-HYm20 | male | none | n/a | XIX |
| Japan | Hyogo | JAP-HYm21 | male | none | n/a | XIX |
| Japan | Hyogo | JAP-HYm23 | male | undetermined | n/a | VI |
| Japan | Hyogo | JAP-HYm24 | male | none | n/a | VI |
| Japan | Hyogo | JAP-HYm25 | male | none | n/a | I |

| | | | | | | |
|---|---|---|---|---|---|---|
| Japan | Hyogo | JAP-HYm26 | male | none | n/a | VI |
| Japan | Hyogo | JAP-HYm27 | male | none | n/a | VI |
| Japan | Hyogo | JAP-HYm28 | male | none | n/a | VI |
| Japan | Hyogo | JAP-HYm29 | male | none | n/a | VI |
| Japan | Hyogo | JAP-HYm30 | male | none | n/a | XIX |
| Japan | Tottori | JAP-TOf01 | female | none | n/a | I |
| Japan | Tottori | JAP-TOf02 | female | none | n/a | I |
| Japan | Tottori | JAP-TOf03 | female | none | n/a | I |
| Japan | Tottori | JAP-TOf04 | female | none | n/a | I |
| Japan | Tottori | JAP-TOf05 | female | none | n/a | VI |
| Japan | Tottori | JAP-TOf06 | female | none | n/a | I |
| Japan | Tottori | JAP-TOf07 | female | none | n/a | VI |
| Japan | Tottori | JAP-TOf08 | female | none | n/a | VI |
| Japan | Tottori | JAP-TOf09 | female | none | n/a | I |
| Japan | Tottori | JAP-TOf10 | female | none | n/a | I |
| Japan | Tottori | JAP-TOf11 | female | none | n/a | VI |
| Japan | Tottori | JAP-TOf12 | female | none | n/a | I |
| Japan | Tottori | JAP-TOf13 | female | none | n/a | VI |
| Japan | Tottori | JAP-TOf14 | female | none | n/a | VI |
| Japan | Tottori | JAP-TOf15 | female | none | n/a | I |
| Japan | Tottori | JAP-TOf16 | female | none | n/a | VI |
| Japan | Tottori | JAP-TOf17 | female | none | n/a | I |
| Japan | Tottori | JAP-TOf18 | female | none | n/a | I |
| Japan | Tottori | JAP-TOf19 | female | none | n/a | I |
| Japan | Tottori | JAP-TOf20 | female | none | n/a | I |
| Japan | Tottori | JAP-TOf21 | female | none | n/a | VI |
| Japan | Tottori | JAP-TOf22 | female | none | n/a | I |
| Japan | Tottori | JAP-TOf23 | female | none | n/a | VI |
| Japan | Tottori | JAP-TOf25 | female | f element | n/a | I |
| Japan | Tottori | JAP-TOf26 | female | none | n/a | I |
| Japan | Tottori | JAP-TOf28 | female | none | n/a | I |
| Japan | Tottori | JAP-TOf29 | female | none | n/a | I |
| Japan | Tottori | JAP-TOf30 | female | f element | n/a | I |
| Japan | Tottori | JAP-TOm01 | male | none | n/a | I |
| Japan | Tottori | JAP-TOm02 | male | none | n/a | VI |
| Japan | Tottori | JAP-TOm04 | male | none | n/a | I |
| Japan | Tottori | JAP-TOm05 | male | none | n/a | I |
| Japan | Tottori | JAP-TOm06 | male | none | n/a | VI |
| Japan | Tottori | JAP-TOm08 | male | none | n/a | I |
| Japan | Tottori | JAP-TOm09 | male | none | n/a | I |
| Japan | Tottori | JAP-TOm10 | male | none | n/a | VI |
| Japan | Tottori | JAP-TOm13 | male | none | n/a | VI |
| Japan | Tottori | JAP-TOm14 | male | none | n/a | VI |
| Japan | Tottori | JAP-TOm16 | male | none | n/a | I |
| Japan | Tottori | JAP-TOm17 | male | none | n/a | VI |
| Japan | Tottori | JAP-TOm21 | male | none | n/a | I |
| Japan | Tottori | JAP-TOm22 | male | none | n/a | VI |
| Japan | Tottori | JAP-TOm23 | male | none | n/a | I |
| Japan | Tottori | JAP-TOm24 | male | none | n/a | I |
| Japan | Tottori | JAP-TOm25 | male | none | n/a | VI |
| Japan | Tottori | JAP-TOm26 | male | none | n/a | I |

| | | | | | | |
|---|---|---|---|---|---|---|
| Japan | Tottori | JAP-TOm28 | male | none | n/a | I |
| Japan | Tottori | JAP-TOm29 | male | none | n/a | VI |
| Japan | Tottori | JAP-TOm30 | male | none | n/a | VI |
| Czech Republic | Prague | CZR-PRf01 | female | f element | n/a | I |
| Czech Republic | Prague | CZR-PRf02 | female | f element | n/a | I |
| Czech Republic | Prague | CZR-PRf03 | female | f element | n/a | I |
| Czech Republic | Prague | CZR-PRf04 | female | f element | n/a | I |
| Czech Republic | Prague | CZR-PRf05 | female | f element | n/a | I |
| Czech Republic | Prague | CZR-PRf06 | female | f element | n/a | I |
| Czech Republic | Prague | CZR-PRf07 | female | f element | n/a | I |
| Czech Republic | Prague | CZR-PRf08 | female | f element | n/a | I |
| Czech Republic | Prague | CZR-PRf09 | female | none | n/a | I |
| Czech Republic | Prague | CZR-PRf10 | female | f element | n/a | I |
| Czech Republic | Prague | CZR-PRf11 | female | f element | n/a | I |
| Czech Republic | Prague | CZR-PRf12 | female | f element | n/a | I |
| Czech Republic | Prague | CZR-PRf13 | female | f element | n/a | I |
| Czech Republic | Prague | CZR-PRf14 | female | f element | n/a | I |
| Czech Republic | Prague | CZR-PRf15 | female | f element | n/a | I |
| Czech Republic | Prague | CZR-PRf16 | female | f element | n/a | I |
| Czech Republic | Prague | CZR-PRf17 | female | f element | n/a | I |
| Czech Republic | Prague | CZR-PRf18 | female | f element | n/a | I |
| Czech Republic | Prague | CZR-PRf19 | female | f element | n/a | I |
| Czech Republic | Prague | CZR-PRf20 | female | f element | n/a | I |
| Czech Republic | Prague | CZR-PRf21 | female | f element | n/a | I |
| Czech Republic | Prague | CZR-PRf22 | female | f element | n/a | I |
| Czech Republic | Prague | CZR-PRf23 | female | f element | n/a | I |
| Czech Republic | Prague | CZR-PRf24 | female | f element | n/a | I |
| Czech Republic | Prague | CZR-PRf25 | female | f element | n/a | I |
| Czech Republic | Prague | CZR-PRf26 | female | f element | n/a | I |
| Czech Republic | Prague | CZR-PRf27 | female | f element | n/a | I |
| Czech Republic | Prague | CZR-PRm01 | male | none | n/a | I |
| Czech Republic | Prague | CZR-PRm02 | male | none | n/a | I |
| Czech Republic | Prague | CZR-PRm03 | male | none | n/a | I |
| Czech Republic | Prague | CZR-PRm04 | male | none | n/a | I |
| Czech Republic | Prague | CZR-PRm05 | male | none | n/a | I |
| Czech Republic | Prague | CZR-PRm06 | male | none | n/a | I |
| Czech Republic | Prague | CZR-PRm07 | male | none | n/a | I |
| Czech Republic | Prague | CZR-PRm08 | male | none | n/a | I |
| Czech Republic | Prague | CZR-PRm09 | male | none | n/a | I |
| Italy | Pisa | ITA-PIf03 | female | none | n/a | V |
| Italy | Pisa | ITA-PIf04 | female | none | n/a | XIII |
| Italy | Pisa | ITA-PIf05 | female | Wolbachia | wVulC | V |
| Italy | Pisa | ITA-PIf06 | female | none | n/a | XIII |
| Italy | Pisa | ITA-PIf07 | female | none | n/a | XIV |
| Italy | Pisa | ITA-PIf08 | female | none | n/a | XIII |
| Italy | Pisa | ITA-PIf09 | female | Wolbachia | wVulC | V |
| Italy | Pisa | ITA-PIf10 | female | Wolbachia | wVulC | V |
| Italy | Pisa | ITA-PIf11 | female | none | n/a | V |
| Italy | Pisa | ITA-PIf12 | female | none | n/a | XIV |
| Italy | Pisa | ITA-PIf13 | female | none | n/a | XV |
| Italy | Pisa | ITA-PIf14 | female | none | n/a | XV |

| | | | | | | |
|---|---|---|---|---|---|---|
| Italy | Pisa | ITA-PIf15 | female | none | n/a | XIV |
| Italy | Pisa | ITA-PIm01 | male | none | n/a | XIII |
| Italy | Pisa | ITA-PIm02 | male | none | n/a | XIII |
| Italy | Pisa | ITA-PIm03 | male | none | n/a | XVI |
| Italy | Pisa | ITA-PIm04 | male | none | n/a | XVII |
| Italy | Pisa | ITA-PIm05 | male | none | n/a | XIV |
| Italy | Pisa | ITA-PIm06 | male | none | n/a | XIV |
| Italy | Pisa | ITA-PIm07 | male | none | n/a | V |
| Italy | Pisa | ITA-PIm08 | male | none | n/a | XIV |
| Italy | Pisa | ITA-PIm09 | male | none | n/a | XV |
| Italy | Pisa | ITA-PIm10 | male | none | n/a | V |
| Italy | Pisa | ITA-PIm11 | male | none | n/a | XVIII |
| Italy | Pisa | ITA-PIm12 | male | none | n/a | XV |
| Italy | Pisa | ITA-PIm13 | male | none | n/a | V |
| Italy | Pisa | ITA-PIm14 | male | none | n/a | XIV |
| Italy | Pisa | ITA-PIm15 | male | none | n/a | V |
| Romania | Bucharest | ROM-BUf01 | female | none | n/a | XX |
| Romania | Bucharest | ROM-BUf02 | female | none | n/a | XX |
| Romania | Bucharest | ROM-BUf03 | female | none | n/a | XX |
| Romania | Bucharest | ROM-BUf04 | female | none | n/a | XX |
| Romania | Bucharest | ROM-BUf05 | female | none | n/a | XXI |
| Romania | Bucharest | ROM-BUf06 | female | none | n/a | XXI |
| Romania | Bucharest | ROM-BUf07 | female | none | n/a | XX |
| Romania | Bucharest | ROM-BUf08 | female | none | n/a | XXII |
| Romania | Bucharest | ROM-BUm01 | male | none | n/a | XXIII |
| Romania | Bucharest | ROM-BUm02 | male | none | n/a | XX |
| Romania | Bucharest | ROM-BUm03 | male | none | n/a | XX |
| Romania | Bucharest | ROM-BUm04 | male | none | n/a | XXIII |
| Romania | Bucharest | ROM-BUm05 | male | none | n/a | XXIII |
| Romania | Bucharest | ROM-BUm06 | male | none | n/a | XX |
| Romania | Bucharest | ROM-BUm07 | male | none | n/a | XX |
| Romania | Bucharest | ROM-BUm08 | male | none | n/a | XXIII |
| Romania | Bucharest | ROM-BUm09 | male | none | n/a | XX |
| Germany | Göttingen | GER-GOf01 | female | f element | n/a | I |
| Germany | Göttingen | GER-GOf02 | female | f element | n/a | I |
| Germany | Göttingen | GER-GOf03 | female | none | n/a | I |
| Germany | Göttingen | GER-GOf04 | female | none | n/a | I |
| Germany | Göttingen | GER-GOf05 | female | Wolbachia | wVulM | II |
| Germany | Göttingen | GER-GOf06 | female | Wolbachia | wVulM | II |
| Germany | Göttingen | GER-GOf07 | female | none | n/a | I |
| Germany | Göttingen | GER-GOf08 | female | none | n/a | I |
| Germany | Göttingen | GER-GOf09 | female | none | n/a | I |
| Germany | Göttingen | GER-GOf10 | female | none | n/a | I |
| Germany | Göttingen | GER-GOf11 | female | none | n/a | I |
| Germany | Göttingen | GER-GOf12 | female | undetermined | n/a | I |
| Germany | Göttingen | GER-GOf13 | female | none | n/a | I |
| Germany | Göttingen | GER-GOf14 | female | none | n/a | I |
| Germany | Göttingen | GER-GOf15 | female | f element | n/a | I |
| Germany | Göttingen | GER-GOf16 | female | none | n/a | I |
| Germany | Göttingen | GER-GOf18 | female | none | n/a | I |
| Germany | Göttingen | GER-GOm02 | male | none | n/a | I |

| | | | | | | |
|---|---|---|---|---|---|---|
| Germany | Göttingen | GER-GOm03 | male | none | n/a | I |
| Germany | Göttingen | GER-GOm05 | male | undetermined | n/a | I |
| Germany | Göttingen | GER-GOm06 | male | undetermined | n/a | I |
| Germany | Göttingen | GER-GOm07 | male | undetermined | n/a | I |
| Germany | Göttingen | GER-GOm09 | male | none | n/a | I |
| Germany | Göttingen | GER-GOm11 | male | undetermined | n/a | I |
| Croatia | Lastovo | CRO-LAf01 | female | none | n/a | IX |
| Croatia | Lastovo | CRO-LAf02 | female | none | n/a | IX |
| Croatia | Lastovo | CRO-LAf03 | female | none | n/a | IX |
| Croatia | Lastovo | CRO-LAf04 | female | none | n/a | IX |
| Croatia | Lastovo | CRO-LAf05 | female | none | n/a | IX |
| Croatia | Lastovo | CRO-LAf06 | female | none | n/a | IX |
| Croatia | Lastovo | CRO-LAf07 | female | none | n/a | IX |
| Croatia | Lastovo | CRO-LAf08 | female | none | n/a | IX |
| Croatia | Lastovo | CRO-LAf09 | female | none | n/a | IX |
| Croatia | Lastovo | CRO-LAf10 | female | none | n/a | IX |
| Croatia | Lastovo | CRO-LAf11 | female | none | n/a | IX |
| Croatia | Lastovo | CRO-LAf12 | female | none | n/a | IX |
| Croatia | Lastovo | CRO-LAf13 | female | none | n/a | IX |
| Croatia | Lastovo | CRO-LAf14 | female | none | n/a | IX |
| Croatia | Lastovo | CRO-LAf15 | female | none | n/a | IX |
| Croatia | Lastovo | CRO-LAf16 | female | none | n/a | IX |
| Croatia | Lastovo | CRO-LAf17 | female | none | n/a | IX |
| Croatia | Lastovo | CRO-LAf18 | female | none | n/a | IX |
| Croatia | Lastovo | CRO-LAf19 | female | none | n/a | IX |
| Croatia | Lastovo | CRO-LAf20 | female | none | n/a | IX |
| Croatia | Lastovo | CRO-LAf21 | female | none | n/a | IX |
| Croatia | Lastovo | CRO-LAf22 | female | none | n/a | IX |
| Croatia | Lastovo | CRO-LAf23 | female | none | n/a | IX |
| Croatia | Lastovo | CRO-LAf24 | female | none | n/a | IX |
| Croatia | Lastovo | CRO-LAm01 | male | none | n/a | IX |
| Croatia | Lastovo | CRO-LAm02 | male | none | n/a | IX |
| Croatia | Lastovo | CRO-LAm03 | male | none | n/a | IX |
| Croatia | Lastovo | CRO-LAm04 | male | none | n/a | IX |
| Croatia | Lastovo | CRO-LAm05 | male | none | n/a | IX |
| Croatia | Lastovo | CRO-LAm06 | male | none | n/a | IX |
| Croatia | Lastovo | CRO-LAm07 | male | none | n/a | IX |
| Croatia | Lastovo | CRO-LAm08 | male | none | n/a | IX |
| Croatia | Lastovo | CRO-LAm09 | male | none | n/a | IX |
| Croatia | Lastovo | CRO-LAm10 | male | none | n/a | IX |
| Croatia | Lastovo | CRO-LAm11 | male | none | n/a | IX |
| Croatia | Lastovo | CRO-LAm12 | male | none | n/a | IX |
| Croatia | Lastovo | CRO-LAm13 | male | none | n/a | IX |
| Croatia | Lastovo | CRO-LAm14 | male | none | n/a | IX |
| Croatia | Lastovo | CRO-LAm15 | male | none | n/a | IX |
| Croatia | Lastovo | CRO-LAm16 | male | none | n/a | IX |
| Croatia | Lastovo | CRO-LAm17 | male | none | n/a | IX |
| Croatia | Lastovo | CRO-LAm18 | male | none | n/a | IX |
| Croatia | Lastovo | CRO-LAm19 | male | none | n/a | IX |
| Croatia | Lastovo | CRO-LAm20 | male | none | n/a | IX |
| Croatia | Lastovo | CRO-LAm21 | male | none | n/a | IX |

| | | | | | | |
|---|---|---|---|---|---|---|
| Croatia | Lastovo | CRO-LAm22 | male | none | n/a | IX |
| Croatia | Lastovo | CRO-LAm23 | male | none | n/a | IX |
| Croatia | Lastovo | CRO-LAm24 | male | none | n/a | IX |
| Croatia | Lastovo | CRO-LAm25 | male | none | n/a | X |
| Croatia | Lastovo | CRO-LAm26 | male | none | n/a | IX |
| Croatia | Lastovo | CRO-LAm27 | male | none | n/a | IX |
| Croatia | Lastovo | CRO-LAm28 | male | none | n/a | IX |
| Croatia | Lastovo | CRO-LAm29 | male | none | n/a | IX |
| Croatia | Lastovo | CRO-LAm30 | male | none | n/a | IX |
| France | Floirac | FRA-FLf01 | female | Wolbachia | wVulC | V |
| France | Floirac | FRA-FLf02 | female | none | n/a | VI |
| France | Floirac | FRA-FLf03 | female | Wolbachia | wVulC | V |
| France | Floirac | FRA-FLf04 | female | Wolbachia | wVulC | V |
| France | Floirac | FRA-FLf05 | female | Wolbachia | wVulC | V |
| France | Floirac | FRA-FLf06 | female | Wolbachia | wVulC | V |
| France | Floirac | FRA-FLf07 | female | none | n/a | VI |
| France | Floirac | FRA-FLf08 | female | Wolbachia | wVulM | II |
| France | Floirac | FRA-FLf09 | female | Wolbachia | wVulM | II |
| France | Floirac | FRA-FLf10 | female | Wolbachia | wVulC | V |
| France | Floirac | FRA-FLf11 | female | Wolbachia | wVulC | V |
| France | Floirac | FRA-FLf12 | female | Wolbachia | wVulC | V |
| France | Floirac | FRA-FLf13 | female | Wolbachia | wVulM | II |
| France | Floirac | FRA-FLf14 | female | Wolbachia | wVulC | V |
| France | Floirac | FRA-FLf15 | female | f element | n/a | V |
| France | Floirac | FRA-FLf16 | female | none | n/a | VI |
| France | Floirac | FRA-FLf17 | female | Wolbachia | wVulC | V |
| France | Floirac | FRA-FLf18 | female | none | n/a | V |
| France | Floirac | FRA-FLf19 | female | Wolbachia | wVulC | V |
| France | Floirac | FRA-FLf20 | female | Wolbachia | wVulC | V |
| France | Floirac | FRA-FLf21 | female | none | n/a | VI |
| France | Floirac | FRA-FLf22 | female | Wolbachia | wVulM | II |
| France | Floirac | FRA-FLf23 | female | Wolbachia | wVulC | V |
| France | Floirac | FRA-FLf24 | female | Wolbachia | wVulC | V |
| France | Floirac | FRA-FLf25 | female | Wolbachia | wVulC | V |
| France | Floirac | FRA-FLf26 | female | Wolbachia | wVulC | V |
| France | Floirac | FRA-FLf27 | female | Wolbachia | wVulM | II |
| France | Floirac | FRA-FLf28 | female | Wolbachia | wVulC | V |
| France | Floirac | FRA-FLf29 | female | Wolbachia | wVulC | V |
| France | Floirac | FRA-FLf30 | female | Wolbachia | wVulC | V |
| France | Floirac | FRA-FLf31 | female | Wolbachia | wVulC | V |
| France | Floirac | FRA-FLf32 | female | Wolbachia | wVulC | V |
| France | Floirac | FRA-FLf34 | female | f element | n/a | VI |
| France | Floirac | FRA-FLf35 | female | none | n/a | VI |
| France | Floirac | FRA-FLf36 | female | Wolbachia | wVulC | V |
| France | Floirac | FRA-FLf37 | female | Wolbachia | wVulC | V |
| France | Floirac | FRA-FLf38 | female | Wolbachia | wVulC | V |
| France | Floirac | FRA-FLf39 | female | Wolbachia | wVulC | V |
| France | Floirac | FRA-FLf40 | female | none | n/a | VI |
| France | Floirac | FRA-FLf41 | female | none | n/a | V |
| France | Floirac | FRA-FLf42 | female | Wolbachia | wVulC | V |
| France | Floirac | FRA-FLf43 | female | f element | n/a | V |

| | | | | | | |
|---|---|---|---|---|---|---|
| France | Floirac | FRA-FLf44 | female | Wolbachia | wVulM | II |
| France | Floirac | FRA-FLf45 | female | Wolbachia | wVulC | V |
| France | Floirac | FRA-FLf46 | female | Wolbachia | wVulC | V |
| France | Floirac | FRA-FLf47 | female | none | n/a | VI |
| France | Floirac | FRA-FLf48 | female | f element | n/a | VI |
| France | Floirac | FRA-FLf49 | female | Wolbachia | wVulC | V |
| France | Floirac | FRA-FLf50 | female | f element | n/a | VI |
| France | Floirac | FRA-FLf51 | female | Wolbachia | wVulC | V |
| France | Floirac | FRA-FLf52 | female | none | n/a | V |
| France | Floirac | FRA-FLf53 | female | Wolbachia | wVulM | II |
| France | Floirac | FRA-FLf54 | female | Wolbachia | wVulC | V |
| France | Floirac | FRA-FLf55 | female | Wolbachia | wVulM | II |
| France | Floirac | FRA-FLf56 | female | none | n/a | V |
| France | Floirac | FRA-FLf57 | female | Wolbachia | wVulC | V |
| France | Floirac | FRA-FLf58 | female | Wolbachia | wVulC | V |
| France | Floirac | FRA-FLf59 | female | Wolbachia | wVulC | XI |
| France | Floirac | FRA-FLf60 | female | f element | n/a | V |
| France | Floirac | FRA-FLf61 | female | Wolbachia | wVulC | V |
| France | Floirac | FRA-FLf62 | female | Wolbachia | wVulC | V |
| France | Floirac | FRA-FLf63 | female | none | n/a | V |
| France | Floirac | FRA-FLf64 | female | none | n/a | VI |
| France | Floirac | FRA-FLf65 | female | none | n/a | VI |
| France | Floirac | FRA-FLf66 | female | none | n/a | V |
| France | Floirac | FRA-FLf67 | female | Wolbachia | wVulC | V |
| France | Floirac | FRA-FLf68 | female | Wolbachia | wVulM | II |
| France | Floirac | FRA-FLf69 | female | Wolbachia | wVulC | V |
| France | Floirac | FRA-FLf70 | female | none | n/a | V |
| France | Floirac | FRA-FLf71 | female | Wolbachia | wVulC | V |
| France | Floirac | FRA-FLf72 | female | none | n/a | V |
| France | Floirac | FRA-FLf73 | female | none | n/a | VI |
| France | Floirac | FRA-FLf74 | female | Wolbachia | wVulC | V |
| France | Floirac | FRA-FLf75 | female | none | n/a | VI |
| France | Floirac | FRA-FLf76 | female | none | n/a | V |
| France | Floirac | FRA-FLf77 | female | none | n/a | V |
| France | Floirac | FRA-FLm01 | male | Wolbachia | wVulC | V |
| France | Floirac | FRA-FLm02 | male | none | n/a | V |
| France | Floirac | FRA-FLm03 | male | none | n/a | VI |
| France | Floirac | FRA-FLm04 | male | none | n/a | V |
| France | Floirac | FRA-FLm05 | male | none | n/a | V |
| France | Floirac | FRA-FLm06 | male | none | n/a | VI |
| France | Floirac | FRA-FLm07 | male | none | n/a | V |
| France | Floirac | FRA-FLm08 | male | none | n/a | VI |
| France | Floirac | FRA-FLm09 | male | none | n/a | VI |
| France | Floirac | FRA-FLm10 | male | none | n/a | II |
| France | Floirac | FRA-FLm11 | male | none | n/a | II |
| France | Floirac | FRA-FLm12 | male | none | n/a | II |
| France | Floirac | FRA-FLm13 | male | none | n/a | VI |
| France | Floirac | FRA-FLm14 | male | undetermined | n/a | V |
| France | Floirac | FRA-FLm15 | male | none | n/a | V |
| France | Floirac | FRA-FLm16 | male | none | n/a | VI |
| France | Floirac | FRA-FLm17 | male | none | n/a | V |

| | | | | | | |
|---|---|---|---|---|---|---|
| France | Floirac | FRA-FLm18 | male | none | n/a | V |
| France | Floirac | FRA-FLm19 | male | undetermined | n/a | V |
| France | Floirac | FRA-FLm20 | male | none | n/a | VI |
| France | Floirac | FRA-FLm21 | male | none | n/a | II |
| France | Floirac | FRA-FLm22 | male | Wolbachia | wVulC | V |
| France | Floirac | FRA-FLm23 | male | none | n/a | V |
| France | Floirac | FRA-FLm24 | male | none | n/a | VI |
| France | Floirac | FRA-FLm25 | male | none | n/a | VI |
| France | Floirac | FRA-FLm26 | male | none | n/a | V |
| France | Floirac | FRA-FLm27 | male | none | n/a | VI |
| France | Floirac | FRA-FLm28 | male | none | n/a | II |
| France | Floirac | FRA-FLm29 | male | none | n/a | V |
| France | Floirac | FRA-FLm30 | male | none | n/a | VI |
| France | Floirac | FRA-FLm31 | male | none | n/a | V |
| France | Floirac | FRA-FLm32 | male | none | n/a | V |
| France | Floirac | FRA-FLm33 | male | none | n/a | V |
| France | Floirac | FRA-FLm34 | male | none | n/a | V |
| France | Floirac | FRA-FLm35 | male | none | n/a | II |
| France | Floirac | FRA-FLm36 | male | none | n/a | V |
| France | Floirac | FRA-FLm37 | male | none | n/a | II |
| France | Floirac | FRA-FLm38 | male | none | n/a | V |
| France | Saint Julien L'Ars | FRA-SJf01 | female | Wolbachia | wVulM | II |
| France | Saint Julien L'Ars | FRA-SJf02 | female | none | n/a | V |
| France | Saint Julien L'Ars | FRA-SJf03 | female | Wolbachia | wVulM | II |
| France | Saint Julien L'Ars | FRA-SJf04 | female | Wolbachia | wVulC | V |
| France | Saint Julien L'Ars | FRA-SJf05 | female | Wolbachia | wVulM | II |
| France | Saint Julien L'Ars | FRA-SJf06 | female | undetermined | n/a | I |
| France | Saint Julien L'Ars | FRA-SJf07 | female | Wolbachia | wVulC | V |
| France | Saint Julien L'Ars | FRA-SJf08 | female | Wolbachia | wVulC | XII |
| France | Saint Julien L'Ars | FRA-SJf09 | female | Wolbachia | wVulC | V |
| France | Saint Julien L'Ars | FRA-SJf10 | female | Wolbachia | wVulC | V |
| France | Saint Julien L'Ars | FRA-SJf11 | female | Wolbachia | wVulC | V |
| France | Saint Julien L'Ars | FRA-SJf12 | female | Wolbachia | wVulC | V |
| France | Saint Julien L'Ars | FRA-SJf13 | female | Wolbachia | wVulC | XII |
| France | Saint Julien L'Ars | FRA-SJf14 | female | Wolbachia | wVulC | V |
| France | Saint Julien L'Ars | FRA-SJf15 | female | Wolbachia | wVulC | V |
| France | Saint Julien L'Ars | FRA-SJf16 | female | Wolbachia | wVulC | XII |
| France | Saint Julien L'Ars | FRA-SJf17 | female | Wolbachia | wVulC | XII |
| France | Saint Julien L'Ars | FRA-SJm01 | male | Wolbachia | wVulM | II |
| France | Saint Julien L'Ars | FRA-SJm02 | male | Wolbachia | wVulM | II |
| France | Saint Julien L'Ars | FRA-SJm03 | male | Wolbachia | not available | II |
| France | Saint Julien L'Ars | FRA-SJm04 | male | none | n/a | I |
| France | Saint Julien L'Ars | FRA-SJm05 | male | Wolbachia | wVulM | II |
| France | Saint Julien L'Ars | FRA-SJm06 | male | none | n/a | I |
| France | Saint Julien L'Ars | FRA-SJm07 | male | none | n/a | V |
| France | Saint Julien L'Ars | FRA-SJm08 | male | none | n/a | II |
| France | Saint Julien L'Ars | FRA-SJm09 | male | none | n/a | I |
| France | Saint Julien L'Ars | FRA-SJm10 | male | none | n/a | I |
| France | Saint Julien L'Ars | FRA-SJm11 | male | none | n/a | V |
| France | Saint Julien L'Ars | FRA-SJm12 | male | Wolbachia | wVulC | V |
| France | Saint Julien L'Ars | FRA-SJm13 | male | none | n/a | XII |

| | | | | | | |
|---|---|---|---|---|---|---|
| France | Saint Julien L'Ars | FRA-SJm14 | male | none | n/a | V |
| The Netherlands | Wageningen | NET-WBf01 | female | none | n/a | VI |
| The Netherlands | Wageningen | NET-WBf03 | female | none | n/a | VI |
| The Netherlands | Wageningen | NET-WBf05 | female | none | n/a | VI |
| The Netherlands | Wageningen | NET-WBf06 | female | none | n/a | VI |
| The Netherlands | Wageningen | NET-WBf08 | female | none | n/a | VI |
| The Netherlands | Wageningen | NET-ABf02 | female | undetermined | n/a | VI |
| The Netherlands | Wageningen | NET-ABf04 | female | none | n/a | II |
| The Netherlands | Wageningen | NET-ADf06 | female | none | n/a | I |
| The Netherlands | Wageningen | NET-WJf02 | female | f element | n/a | VI |
| The Netherlands | Wageningen | NET-ABm02 | male | none | n/a | VI |
| The Netherlands | Wageningen | NET-WJm02 | male | none | n/a | VI |
| France | Beauvoir | BE17f01 | female | undetermined | n/a | I |
| France | Beauvoir | BE17f02 | female | f element | n/a | I |
| France | Beauvoir | BE17f03 | female | none | n/a | I |
| France | Beauvoir | BE17f04 | female | Wolbachia | wVulM | II |
| France | Beauvoir | BE17f05 | female | none | n/a | I |
| France | Beauvoir | BE17f06 | female | none | n/a | I |
| France | Beauvoir | BE17f07 | female | f element | n/a | I |
| France | Beauvoir | BE17f08 | female | f element | n/a | I |
| France | Beauvoir | BE17f09 | female | f element | n/a | I |
| France | Beauvoir | BE17f10 | female | f element | n/a | I |
| France | Beauvoir | BE17f11 | female | none | n/a | I |
| France | Beauvoir | BE17f12 | female | f element | n/a | I |
| France | Beauvoir | BE17f13 | female | none | n/a | I |
| France | Beauvoir | BE17f14 | female | f element | n/a | I |
| France | Beauvoir | BE17f15 | female | none | n/a | I |
| France | Beauvoir | BE17f16 | female | f element | n/a | I |
| France | Beauvoir | BE17f17 | female | none | n/a | I |
| France | Beauvoir | BE17f18 | female | none | n/a | I |
| France | Beauvoir | BE17f19 | female | f element | n/a | I |
| France | Beauvoir | BE17f20 | female | none | n/a | I |
| France | Beauvoir | BE17f21 | female | f element | n/a | I |
| France | Beauvoir | BE17f22 | female | f element | n/a | I |
| France | Beauvoir | BE17f24 | female | f element | n/a | I |
| France | Beauvoir | BE17f25 | female | f element | n/a | I |
| France | Beauvoir | BE17f26 | female | f element | n/a | I |
| France | Beauvoir | BE17m01 | male | none | n/a | I |
| France | Beauvoir | BE17m03 | male | f element | n/a | I |
| France | Beauvoir | BE17m04 | male | none | n/a | I |
| France | Beauvoir | BE17m05 | male | none | n/a | I |
| France | Beauvoir | BE17m07 | male | none | n/a | I |
| France | Beauvoir | BE17m09 | male | none | n/a | I |
| France | Chizé | CH17f01 | female | f element | n/a | III |
| France | Chizé | CH17f02 | female | f element | n/a | II |
| France | Chizé | CH17f03 | female | f element | n/a | III |
| France | Chizé | CH17f05 | female | none | n/a | III |
| France | Chizé | CH17f06 | female | f element | n/a | III |
| France | Chizé | CH17f07 | female | Wolbachia and f element | wVulM | II |
| France | Chizé | CH17f08 | female | none | n/a | III |

| | | | | | | |
|---|---|---|---|---|---|---|
| France | Chizé | CH17f09 | female | f element | n/a | III |
| France | Chizé | CH17f10 | female | f element | n/a | III |
| France | Chizé | CH17f11 | female | f element | n/a | III |
| France | Chizé | CH17f12 | female | Wolbachia and f element | wVulM | II |
| France | Chizé | CH17f13 | female | f element | n/a | III |
| France | Chizé | CH17f14 | female | f element | n/a | III |
| France | Chizé | CH17f15 | female | f element | n/a | III |
| France | Chizé | CH17f16 | female | f element | n/a | III |
| France | Chizé | CH17f17 | female | f element | n/a | III |
| France | Chizé | CH17f18 | female | f element | n/a | III |
| France | Chizé | CH17f19 | female | f element | n/a | III |
| France | Chizé | CH17f20 | female | f element | n/a | III |
| France | Chizé | CH17f21 | female | f element | n/a | III |
| France | Chizé | CH17f22 | female | f element | n/a | III |
| France | Chizé | CH17f23 | female | f element | n/a | III |
| France | Chizé | CH17f24 | female | f element | n/a | II |
| France | Chizé | CH17f25 | female | f element | n/a | II |
| France | Chizé | CH17f26 | female | Wolbachia | wVulM | II |
| France | Chizé | CH17f27 | female | f element | n/a | III |
| France | Chizé | CH17f28 | female | f element | n/a | III |
| France | Chizé | CH17f29 | female | f element | n/a | III |
| France | Chizé | CH17f30 | female | f element | n/a | III |
| France | Chizé | CH17f31 | female | Wolbachia and f element | wVulM | II |
| France | Chizé | CH17f32 | female | f element | n/a | II |
| France | Chizé | CH17f33 | female | f element | n/a | III |
| France | Chizé | CH17f34 | female | f element | n/a | III |
| France | Chizé | CH17f35 | female | f element | n/a | III |
| France | Chizé | CH17f36 | female | f element | n/a | III |
| France | Chizé | CH17f37 | female | f element | n/a | III |
| France | Chizé | CH17f38 | female | f element | n/a | III |
| France | Chizé | CH17f39 | female | f element | n/a | III |
| France | Chizé | CH17f40 | female | none | n/a | III |
| France | Chizé | CH17f44 | female | f element | n/a | III |
| France | Chizé | CH17f45 | female | Wolbachia | wVulM | II |
| France | Chizé | CH17f46 | female | f element | n/a | III |
| France | Chizé | CH17f47 | female | f element | n/a | III |
| France | Chizé | CH17f48 | female | f element | n/a | III |
| France | Chizé | CH17m02 | male | f element | n/a | III |
| France | Chizé | CH17m03 | male | none | n/a | III |
| France | Chizé | CH17m04 | male | f element | n/a | III |
| France | Chizé | CH17m05 | male | f element | n/a | III |
| France | Chizé | CH17m06 | male | f element | n/a | III |
| France | Chizé | CH17m07 | male | none | n/a | III |
| France | Chizé | CH17m08 | male | f element | n/a | II |
| France | Chizé | CH17m09 | male | f element | n/a | IV |
| France | Coulombiers | CO17f01 | female | f element | n/a | V |
| France | Coulombiers | CO17f02 | female | f element | n/a | V |
| France | Coulombiers | CO17f03 | female | f element | n/a | I |
| France | Coulombiers | CO17f04 | female | f element | n/a | I |

| | | | | | | |
|---|---|---|---|---|---|---|
| France | Coulombiers | CO17f05 | female | f element | n/a | I |
| France | Coulombiers | CO17f06 | female | f element | n/a | I |
| France | Coulombiers | CO17f07 | female | none | n/a | I |
| France | Coulombiers | CO17f08 | female | none | n/a | VI |
| France | Coulombiers | CO17f09 | female | none | n/a | VI |
| France | Coulombiers | CO17f10 | female | none | n/a | I |
| France | Coulombiers | CO17f11 | female | f element | n/a | V |
| France | Coulombiers | CO17f12 | female | f element | n/a | VI |
| France | Coulombiers | CO17f13 | female | f element | n/a | I |
| France | Coulombiers | CO17f14 | female | f element | n/a | I |
| France | Coulombiers | CO17f15 | female | f element | n/a | I |
| France | Coulombiers | CO17f16 | female | f element | n/a | I |
| France | Coulombiers | CO17f17 | female | Wolbachia | wVulC | V |
| France | Coulombiers | CO17f18 | female | none | n/a | V |
| France | Coulombiers | CO17f19 | female | f element | n/a | V |
| France | Coulombiers | CO17f20 | female | none | n/a | VII |
| France | Coulombiers | CO17m01 | male | none | n/a | I |
| France | Coulombiers | CO17m02 | male | f element | n/a | V |
| France | Coulombiers | CO17m03 | male | f element | n/a | I |
| France | Coulombiers | CO17m04 | male | none | n/a | VI |
| France | Gript | GR17f01 | female | none | n/a | III |
| France | Gript | GR17f02 | female | f element | n/a | I |
| France | Gript | GR17f03 | female | none | n/a | III |
| France | Gript | GR17f04 | female | none | n/a | III |
| France | Gript | GR17f05 | female | none | n/a | III |
| France | Gript | GR17f06 | female | none | n/a | III |
| France | Gript | GR17f07 | female | none | n/a | III |
| France | Gript | GR17f08 | female | none | n/a | III |
| France | Gript | GR17f09 | female | none | n/a | III |
| France | Gript | GR17f10 | female | Wolbachia | wVulC | V |
| France | Gript | GR17f11 | female | none | n/a | III |
| France | Gript | GR17f12 | female | none | n/a | III |
| France | Gript | GR17f13 | female | none | n/a | III |
| France | Gript | GR17f14 | female | none | n/a | III |
| France | Gript | GR17f15 | female | none | n/a | III |
| France | Gript | GR17f16 | female | none | n/a | III |
| France | Gript | GR17f17 | female | none | n/a | III |
| France | Gript | GR17f18 | female | none | n/a | III |
| France | Gript | GR17f19 | female | Wolbachia | wVulC | V |
| France | Gript | GR17f20 | female | none | n/a | III |
| France | Gript | GR17f21 | female | none | n/a | III |
| France | Gript | GR17f22 | female | none | n/a | III |
| France | Gript | GR17f23 | female | f element | n/a | III |
| France | Gript | GR17f24 | female | none | n/a | III |
| France | Gript | GR17f25 | female | none | n/a | III |
| France | Gript | GR17f26 | female | none | n/a | III |
| France | Gript | GR17f27 | female | none | n/a | III |
| France | Gript | GR17f28 | female | none | n/a | III |
| France | Gript | GR17f29 | female | none | n/a | III |
| France | Gript | GR17f30 | female | none | n/a | III |
| France | Gript | GR17m01 | male | none | n/a | III |

| | | | | | | |
|---|---|---|---|---|---|---|
| France | Gript | GR17m02 | male | none | n/a | III |
| France | Gript | GR17m03 | male | none | n/a | III |
| France | Gript | GR17m04 | male | none | n/a | III |
| France | Gript | GR17m05 | male | none | n/a | III |
| France | Gript | GR17m06 | male | none | n/a | III |
| France | Gript | GR17m08 | male | none | n/a | III |
| France | Gript | GR17m09 | male | none | n/a | III |
| France | Gript | GR17m10 | male | none | n/a | III |
| France | Gript | GR17m11 | male | none | n/a | III |
| France | Gript | GR17m12 | male | none | n/a | III |
| France | Gript | GR17m13 | male | none | n/a | III |
| France | Gript | GR17m14 | male | none | n/a | III |
| France | Gript | GR17m15 | male | none | n/a | III |
| France | Gript | GR17m16 | male | none | n/a | III |
| France | La Crèche | LC17f01 | female | none | n/a | I |
| France | La Crèche | LC17f02 | female | none | n/a | VIII |
| France | La Crèche | LC17f03 | female | none | n/a | VIII |
| France | La Crèche | LC17f04 | female | none | n/a | I |
| France | La Crèche | LC17f05 | female | none | n/a | VIII |
| France | La Crèche | LC17f06 | female | none | n/a | I |
| France | La Crèche | LC17f07 | female | f element | n/a | I |
| France | La Crèche | LC17f08 | female | f element | n/a | I |
| France | La Crèche | LC17f10 | female | f element | n/a | I |
| France | La Crèche | LC17f11 | female | none | n/a | I |
| France | La Crèche | LC17f12 | female | none | n/a | I |
| France | La Crèche | LC17f13 | female | f element | n/a | I |
| France | La Crèche | LC17f15 | female | none | n/a | I |
| France | La Crèche | LC17f17 | female | none | n/a | I |
| France | La Crèche | LC17f18 | female | none | n/a | I |
| France | La Crèche | LC17f20 | female | none | n/a | I |
| France | La Crèche | LC17f21 | female | f element | n/a | I |
| France | La Crèche | LC17f22 | female | none | n/a | I |
| France | La Crèche | LC17f23 | female | none | n/a | I |
| France | La Crèche | LC17f25 | female | f element | n/a | I |
| France | La Crèche | LC17f26 | female | f element | n/a | I |
| France | La Crèche | LC17f27 | female | f element | n/a | I |
| France | La Crèche | LC17f28 | female | f element | n/a | I |
| France | La Crèche | LC17f29 | female | none | n/a | I |
| France | La Crèche | LC17f30 | female | f element | n/a | I |
| France | La Crèche | LC17f31 | female | none | n/a | I |
| France | La Crèche | LC17f33 | female | f element | n/a | I |
| France | La Crèche | LC17f35 | female | Wolbachia | wVulC | V |
| France | La Crèche | LC17f36 | female | none | n/a | I |
| France | La Crèche | LC17f37 | female | none | n/a | VIII |
| France | La Crèche | LC17f38 | female | none | n/a | I |
| France | La Crèche | LC17f40 | female | none | n/a | I |
| France | La Crèche | LC17f41 | female | none | n/a | I |
| France | La Crèche | LC17f42 | female | f element | n/a | I |
| France | La Crèche | LC17f43 | female | none | n/a | I |
| France | La Crèche | LC17f44 | female | none | n/a | I |
| France | La Crèche | LC17f45 | female | f element | n/a | I |

| | | | | | | |
|---|---|---|---|---|---|---|
| France | La Crèche | LC17m01 | male | none | n/a | I |
| France | La Crèche | LC17m02 | male | f element | n/a | I |
| France | La Crèche | LC17m04 | male | none | n/a | I |
| France | La Crèche | LC17m05 | male | none | n/a | I |
| France | La Crèche | LC17m06 | male | none | n/a | I |
| France | La Crèche | LC17m07 | male | none | n/a | I |
| France | La Crèche | LC17m08 | male | none | n/a | I |
| France | La Crèche | LC17m09 | male | f element | n/a | I |
| France | La Crèche | LC17m10 | male | none | n/a | I |
| France | La Crèche | LC17m11 | male | none | n/a | I |
| France | La Crèche | LC17m14 | male | none | n/a | not available |
| France | La Crèche | LC17m15 | male | none | n/a | not available |
| France | La Crèche | LC17m16 | male | none | n/a | I |
| France | La Crèche | LC17m17 | male | none | n/a | not available |
| France | La Crèche | LC17m18 | male | none | n/a | VIII |
| France | La Crèche | LC17m19 | male | none | n/a | I |
| France | La Crèche | LC17m20 | male | none | n/a | I |
| France | La Crèche | LC17m21 | male | none | n/a | I |
| France | La Crèche | LC17m22 | male | none | n/a | I |
| France | La Crèche | LC17m23 | male | none | n/a | not available |
| France | La Crèche | LC17m24 | male | none | n/a | not available |
| France | Poitiers | PO15f01 | female | Wolbachia | wVulP | VII |
| France | Poitiers | PO15f02 | female | Wolbachia | not available | VII |
| France | Poitiers | PO15f03 | female | none | n/a | VII |
| France | Poitiers | PO15f04 | female | none | n/a | VII |
| France | Poitiers | PO15f05 | female | none | n/a | VII |
| France | Poitiers | PO15f06 | female | Wolbachia | wVulP | VII |
| France | Poitiers | PO15f07 | female | none | n/a | VII |
| France | Poitiers | PO15f09 | female | Wolbachia | wVulP | VII |
| France | Poitiers | PO15f10 | female | none | n/a | VII |
| France | Poitiers | PO15f11 | female | none | n/a | VII |
| France | Poitiers | PO15f12 | female | Wolbachia | not available | VII |
| France | Poitiers | PO15f13 | female | none | n/a | VII |
| France | Poitiers | PO15f14 | female | f element | n/a | V |
| France | Poitiers | PO15f15 | female | Wolbachia | not available | VII |
| France | Poitiers | PO15f16 | female | none | n/a | VII |
| France | Poitiers | PO15f17 | female | none | n/a | VII |
| France | Poitiers | PO15f18 | female | Wolbachia | wVulP | VII |
| France | Poitiers | PO15f19 | female | none | n/a | V |
| France | Poitiers | PO15f20 | female | Wolbachia | not available | VII |
| France | Poitiers | PO15m01 | male | none | n/a | VII |
| France | Poitiers | PO15m03 | male | none | n/a | VII |
| France | Poitiers | PO15m04 | male | none | n/a | VII |
| France | Poitiers | PO15m06 | male | none | n/a | VII |

**Table S3. Relative dose of the *Wolbachia* marker for the samples tested in quantitative PCR.** For each assay, the cycle threshold (CT) of the reference gene (tleu) and of the target (z12), the sex of the individual and the relative dose of *Wolbachia* is given.

| Individual ID | CT-z12 | CT-tleu | Relative dose | Sex |
|---|---|---|---|---|
| FRA-FLf-08 | 25.5824435 | 25.6882256 | 1.076 | Female |
| FRA-FLf-09 | 23.6277411 | 23.5971254 | 0.979 | Female |
| FRA-FLf-10 | 24.3705659 | 24.7374311 | 1.29 | Female |
| FRA-FLf-11 | 24.5537862 | 25.7105929 | 2.23 | Female |
| FRA-FLf-12 | 22.3757942 | 22.7804067 | 1.324 | Female |
| FRA-FLm-01 | 21.2115138 | 22.6370199 | 2.686 | Male |
| FRA-FLm-22 | 24.3622963 | 22.3685658 | 0.2511 | Male |
| FRA-SJf-01 | 20.7678607 | 21.8196709 | 2.073 | Female |
| FRA-SJf-03 | 22.5410224 | 22.8779862 | 1.263 | Female |
| FRA-SJf-04 | 21.9990693 | 23.8393105 | 3.581 | Female |
| FRA-SJf-05 | 23.3765567 | 23.4413002 | 1.046 | Female |
| FRA-SJf-07 | 21.8612162 | 22.9316304 | 2.1 | Female |
| FRA-SJf-08 | 21.7508822 | 23.7412059 | 3.973 | Female |
| FRA-SJf-09 | 21.7716541 | 22.9018341 | 2.189 | Female |
| FRA-SJf-10 | 21.6368685 | 22.8778059 | 2.364 | Female |
| FRA-SJf-11 | 21.4285492 | 22.8088711 | 2.603 | Female |
| FRA-SJf-12 | 22.6035473 | 24.4015473 | 3.477 | Female |
| FRA-SJf-13 | 20.9178346 | 22.5161113 | 3.028 | Female |
| FRA-SJf-14 | 20.2761774 | 22.2631569 | 3.964 | Female |
| FRA-SJf-15 | 21.9448917 | 23.4763178 | 2.891 | Female |
| FRA-SJf-16 | 20.626825 | 22.5059827 | 3.679 | Female |
| FRA-SJf-17 | 19.8270246 | 21.6514128 | 3.542 | Female |
| FRA-SJm-01 | 26.799457 | 22.7215987 | 5.92E-02 | Male |
| FRA-SJm-02 | 23.8809819 | 20.8547259 | 0.1227 | Male |
| FRA-SJm-03 | 28.4457355 | 23.6275762 | 3.54E-02 | Male |
| FRA-SJm-05 | 25.6538476 | 21.1256958 | 4.33E-02 | Male |
| FRA-SJm-12 | 27.2076295 | 22.5279936 | 3.90E-02 | Male |

**Table S4. GenBank accession numbers of 23 mitochondrial haplotypes from the 647 *Armadillidium vulgare* individuals from 16 populations included in this study.**

| Mitochondrial haplotype | GenBank accession number |
|---|---|
| I | OP430857 |
| II | OP430858 |
| III | OP430859 |
| IV | OP430860 |
| V | OP430861 |
| VI | OP430862 |
| VII | OP430863 |
| VIII | OP430864 |
| IX | OP430865 |
| X | OP430866 |
| XI | OP430867 |
| XII | OP430868 |
| XIII | OP430869 |
| XIV | OP430870 |
| XV | OP430871 |
| XVI | OP430872 |
| XVII | OP430873 |
| XVIII | OP430874 |
| XIX | OP430875 |
| XX | OP430876 |
| XXI | OP430877 |
| XXII | OP430878 |
| XXIII | OP430879 |
